# Clustering conformations from molecular dynamics simulations by gradient flow segmentation of projected free energy landscapes

**DOI:** 10.64898/2026.11.25.769211

**Authors:** Mert Gur, Bogdan Kochetov, Mert Golcuk, Ethan M. Bintrim, Fareeda E. Abu-Juam, Shikhar Uttam

## Abstract

Molecular dynamics (MD) simulations sample a continuum of conformations whose populations can reveal distinct protein structural states and substates as well as different ligand binding states. Determining which conformations belong to each state remains a central challenge in interpreting protein structural dynamics and ligand binding. Building on our framework for constructing free-energy surfaces from principal component projections of MD trajectories, we treat these surfaces as images. We adapt a method originally developed to segment cell nuclei in three-dimensional microscopy images and use the basin structure of the free-energy surface as the basis for clustering. We follow gradient flow toward free-energy minima and group neighboring regions by comparing the free energy along their shared boundaries with that within each region. When both regions contain resolved minima, we also test how far apart their cores are relative to their internal variation. Mapping the regions associated with the minima back to the original trajectories recovers the conformational ensemble in each state or substate and allows us to select a representative conformation directly from the simulation. We applied the method to conformational changes in the SARS-CoV-2 spike protein and the leucine transporter LeuT, and to the binding conformations sampled by a small-molecule drug interacting with the PP2A scaffold PR65. By combining statistical mechanics with computer vision, this approach provides a structural interpretation of how sampled conformational populations are organized.

## INTRODUCTION

All-atom molecular dynamics (MD) simulations provide time-resolved descriptions of biomolecular structure and motion that complement experimental measurements.^1^ Their biological interpretation depends on identifying the conformations visited, the structural changes connecting them, and representative structures for the sampled ensembles. A trajectory can sample a broad, unevenly populated distribution of conformations across several structural states. An analysis therefore needs to group the sampled conformations by structural state and link each conformation to its corresponding frame in the original trajectory.

Principal component analysis (PCA) provides a reduced representation of correlated Cartesian fluctuations.^2^ Projecting coordinate frames onto a few principal components can expose densely sampled regions and the less populated areas between them. Such representations have helped characterize collective motions and conformational landscapes.^3^ For each populated bin, the projected free energy is estimated as G = −kBT ln(P) + C, where P is the fraction of trajectory frames in that bin and C sets the energy reference. More frequently sampled regions lie lower on the free-energy landscape. Following the landscape downhill toward local minima identifies the regions occupied by distinct conformational ensembles.

Gradient flow tracking was developed for segmenting touching nuclei in 3D microscopy images by tracking voxels along a diffused gradient field into attraction basins, which are grouped into regions representing individual nuclei.^4^ We adapt this principle to molecular trajectory analysis. In microscopy, an image region often corresponds to an independently identifiable object, such as a nucleus. In a projected free-energy landscape, a segmented region defines an ensemble of sampled conformations whose interpretation depends on the coordinate representation, sampling, and resolution. In both settings, gradient flow delineates regions of attraction and the boundaries between them. On the projected free-energy landscape, these boundaries can be used to locate candidate crossings. For neighboring basins, the free-energy difference between the lowest pass across their shared boundary and the minimum of each basin gives the apparent barrier to crossing from that basin.

Here we introduce a new method that uses the basin structure of projected free-energy surfaces to cluster conformations sampled in MD simulations. The method adapts the gradient flow tracking approach to track gradient flow toward local free-energy minima and groups neighboring regions by comparing the free energy along their shared boundaries with that within each region. A core separation criterion preserves distinct wells in the final grouping. Mapping the resulting basins back to the original trajectory identifies the conformations in each ensemble. The method requires no prescribed basin count. It estimates apparent free-energy barriers between neighboring basins and selects representative conformations directly from the sampled frames. We demonstrate the analysis on SARS-CoV-2 spike trajectories related to closed-to-open motion^5^, LeuT trajectories spanning several starting structures and substrate conditions^6^, and simulations of ATUX-8385 bound to the PP2A scaffold PR65.^7,8^

## METHODOLOGY

We present a gradient flow clustering method that converts MD trajectories into maps of conformational states and substates by delineating basins on projected free-energy surfaces. For each state or substate, the method identifies every assigned frame by its source trajectory and frame index, allowing the full conformational ensemble to be recovered from the original simulations (steps 1-4). The method then locates crossings between neighboring basins and estimates the apparent free-energy barrier from each basin minimum (step 5). Representative conformations for each basin and crossing are selected directly from the sampled frames (step 6). Figure 1 illustrates the method and its structural output for LeuT. Figure 2 shows the intermediate segmentation steps.

**Figure 1.**
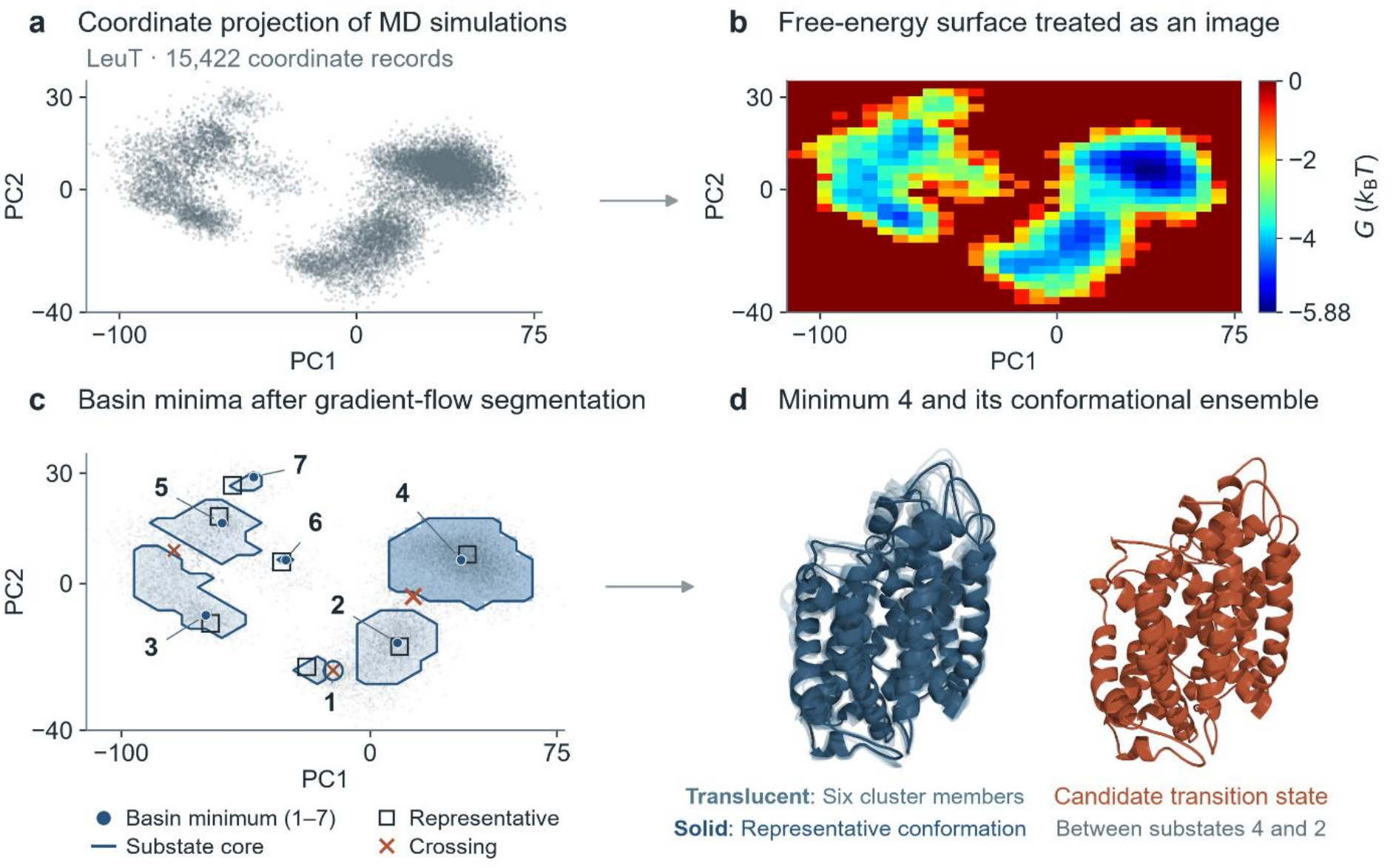
Clustering MD simulation trajectories by gradient flow on projected free-energy surfaces. **(a)** LeuT trajectory frames projected onto the first two principal components (PC1 and PC2). **(b)** Projected free-energy surface treated as an image for segmentation. Blue indicates low free energy. Red marks the zero reference and the unsampled background. **(c)** Basins identified by gradient-flow segmentation. Numbered circles label each basin at its minimum, outlines show basin cores, open squares mark representative conformations, and crosses indicate interbasin crossings. **(d)** Six members of the basin 4 ensemble (translucent blue) are superimposed on its representative conformation (solid blue). The rust-colored conformation represents the candidate transition state between substates 4 and 2.

**Figure 2.**
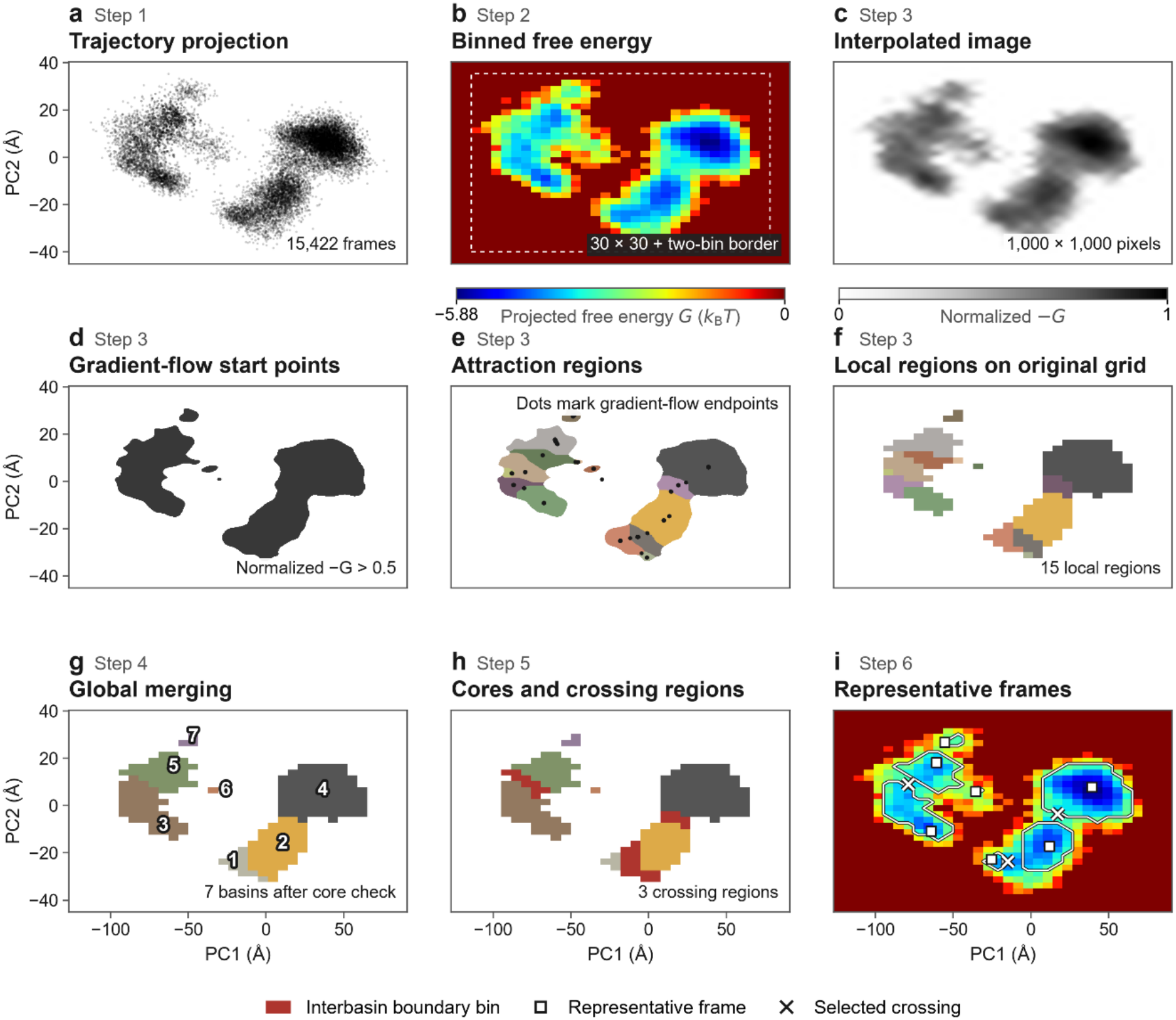
Stages of gradient flow clustering for LeuT. Panel labels identify the corresponding Methods steps. (a) Projection of 15,422 frames onto PC1 and PC2. (b) Free energy on the 30 × 30 grid, a grid size also selected by the automatic rule using the number of sampled frames and the normalized spread along each PC. The dashed line bounds the unpadded grid. (c) Normalized −G image interpolated to 1,000 × 1,000 pixels. (d) Pixels with intensity above 0.5 seed the gradient paths. (e) Colors show attraction regions after small-region reassignment, and dots mark gradient-path endpoints. (f) Transfer of the labels back to the original grid yields 15 local regions. (g) Global merging yields seven basins. (h) Red bins identify three connected interbasin boundary regions. (i) White outlines trace the cores, squares locate representative frames and crosses mark selected crossing bins.

### Method overview

#### Step 1. Molecular representation and projection onto principal components

The choice of atoms and coordinate frame determines which molecular motions are captured in the projection. For a protein undergoing a domain rearrangement, expressing backbone coordinates relative to one domain reveals the motion of the other. For a ligand moving within a binding site, expressing its coordinates relative to the receptor preserves changes in binding position and orientation. In both examples, each trajectory frame is rotated and translated as a whole during trajectory preparation so that the atoms of the chosen domain or receptor best match their positions in a common reference structure. The protein or ligand coordinates used for PCA are then extracted from these aligned frames.

For a selection of N atoms, the configuration vector ***R*** contains the 3*N* Cartesian coordinates of a conformation. Following our previous analyses [3,6], the covariance of the fluctuations about the mean structure is described by the matrix

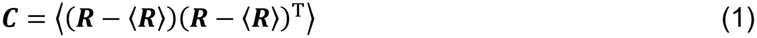

where the brackets denote an equally weighted average over the conformations used to define the principal axes, ⟨***R***⟩ is their mean structure, and the superscript T denotes transpose. Principal component analysis [2] decomposes this matrix as

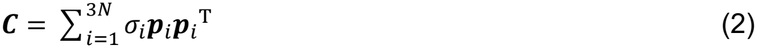

Each eigenvector ***p***_*i*_ gives a direction of conformational change, and its eigenvalue *σ*_*i*_ gives the variance along that direction. Ordering these variances from largest to smallest defines PC1 and PC2 as the two axes along which the sampled conformations vary most. Each eigenvector’s sign is fixed so that its largest-magnitude component is positive, which makes the orientation of the map reproducible.

The position of a conformation along axis *i* is the projection of its displacement from the mean structure onto ***p***_*i*_, given by ***p***_*i*_^T^(***R*** − ⟨***R***⟩). For a set of trajectories, all conformations are projected relative to the same mean structure and onto the same two axes. Each point in the resulting PC1– PC2 map retains its source trajectory and frame number (Fig. 1a).

#### Step 2. Construction of the projected free-energy surface

The distribution of conformations on the PC1–PC2 plane provides an estimate of the projected free-energy surface [3,6]. The plane is divided into equal-area bins. The occupancy *f*_*s*_ of bin *s* is the fraction of sampled conformations that fall within it. For each occupied bin, the free energy is

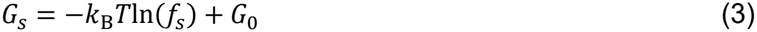

where *k*_B_ is the Boltzmann constant, *T* is the absolute temperature, and *G*_0_ sets the energy reference. The least-populated occupied bin defines the zero of free energy in Figure 1b. *G*_0_ is set to *k*_B_*T*ln(*f*_*min*_), where *f*_*min*_ is the smallest nonzero bin occupancy. More frequently sampled regions therefore lie at negative values of *G* (Figure 1b). Empty bins have no finite free-energy estimate; for segmentation, their values are set to zero in the image. The resulting surface describes the sampled distribution and has an equilibrium interpretation when the bin occupancies represent equilibrium populations.

Users can specify the number of bins along each axis or provide the bin edges. By default, the script selects the number of bins along each PC from the number of projected frames and the relative spread of conformations along that axis. For each PC, its normalized spread, *q*, is the sample standard deviation divided by the 95th percentile of the absolute distances from its median. For *M* projected frames, the target total number of bins is *K* = 65 × log_2_(*M*), where 65 was calibrated using the manually selected LeuT and spike grids. The numbers of bins along PC1 and PC2 are 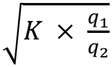 and 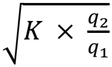, respectively, each rounded to the nearest integer. The grid spans the full range of projected conformations along each axis. By default, two empty bins are added on each side.

#### Step 3. Gradient flow toward local free-energy minima

To delineate the regions surrounding local free-energy minima, the method traces paths toward increasing intensity on the normalized −G image. Cubic B-spline interpolation provides a finer grid on which to estimate the local gradient. If the longer axis contains fewer than 1,000 pixels, both axes are scaled by the same factor to bring the longer axis to approximately 1,000 pixels, the default target. The gradient of the interpolated image is estimated by forward differences and then diffused with the elastic gradient-diffusion scheme of^4^ for 21 explicit iterations (Lamé coefficients μ = 0.05 and λ = 0.01).

Paths start at positions on the interpolated grid where the normalized −G image exceeds a user-defined threshold (default 0.5). The normalization sets empty bins to 0 and the deepest bin to 1, so the threshold is a fraction of the depth of the global minimum below the G = 0 reference. For a nonzero smoothed gradient v(x) at grid position x, the proposed next position x′ is given by the gradient flow rule.^4^

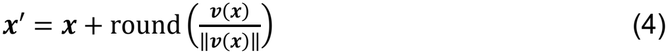

Rounding each component of the unit vector to the nearest integer gives the step to a neighboring pixel. A path stops when it no longer advances, when the next step would leave the image, or when turning angle between the smoothed gradient vectors at the current and proposed next grid positions is 90° or greater.

The original histogram bins set the resolution at which conformational populations are measured. Endpoints within a Euclidean distance of d pixels on the interpolated grid are linked, where d is the interpolation factor rounded to the nearest integer (approximately one original bin spacing), with a lower bound of five pixels. The paths leading to each connected group of endpoints define a local region.

To limit fragmentation, we set the minimum region area to d^2^/3, rounded down to a whole number of pixels, with a lower bound of 25 pixels on the interpolated grid. In the reported analyses, this cutoff corresponded to approximately one-third of the area of an original histogram bin. Regions smaller than this cutoff are reassigned to the most frequent label on their perimeter, excluding labels of other small regions. A small region is left unassigned if the most frequent value on its perimeter is the unlabeled background, or if all labels on its perimeter belong to other small regions. Nearest-neighbor interpolation then transfers the local region labels to the original free-energy grid. Disconnected regions receive separate labels, counting bins that share an edge or a corner as connected.

#### Step 4. Basin delineation and assignment of conformational ensembles

Two neighboring regions are eligible for merging if the mean intensity of the normalized −G image (where higher intensity corresponds to lower free energy) along their shared boundary exceeds the mean intensity over all bins of at least one of the two regions. For occupied bins, this is equivalent to a lower mean free energy along the boundary than within at least one region. These averages are calculated on the original grid, with equal weight given to each bin. The boundary average includes bins on both sides of the shared boundary.

For each eligible pair, we define the current core of each region by excluding bins at its shared boundaries. A core contains a resolved local minimum when a populated bin and its eight immediate neighbors are populated and all belong to that core, and the central free energy is no higher than any of its neighbors and strictly lower than at least one. We evaluate this condition on the original free-energy grid.

When both cores contain a resolved minimum, we calculate the Mahalanobis distance D between the means of their frame coordinates in PC1–PC2. We use the arithmetic mean of the two within-core sample covariance matrices to scale the difference between the means. This accounts for correlation between PCs and gives the two cores equal weight. The distance is dimensionless and invariant to rescaling the PC axes for fixed bin assignments. We keep the regions separate if D exceeds 2. We evaluate the covariance inverse in directions with nonzero variance. Separation along a zero-variance direction prevents merging. Core membership and distance are updated after each merge.

The algorithm begins with the largest unprocessed region, merges the first neighbor in label order that meets the merging criteria, and updates the region labels after each merge. A region is marked as processed when none of its neighbors meets the merging criterion, and the next largest unprocessed region is then examined. A processed region is not examined again as a starting region, but it can still be merged into a neighboring region whose mean intensity changes through later merges. After merging, a region consisting of a single bin takes the most frequent label among its assigned neighbors. Basins are relabeled using edge or corner connectivity. If all bins of a basin lie on shared boundaries and it borders only one neighbor, it is merged into that neighbor and the labels are updated. This step is repeated until no such basin remains, and the number of basins is determined by the resulting partition.

Each frame inherits the basin label of its bin in the original projection. All frames assigned to a basin constitute the conformational ensemble of that state or substate, even when they come from different trajectories or from separate visits to the same basin. Frames in unlabeled bins remain unassigned. Within an ensemble, frames that occupy a shared interbasin boundary retain their basin membership. The remaining frames form the basin core. Figure 1c outlines these cores on the projected surface.

For each basin, the method returns a list of all member frames and identifies those belonging to its core. Each entry gives the source trajectory and frame number needed to retrieve the corresponding coordinates for structural analysis.

#### Step 5. Interbasin crossings and apparent free-energy barriers

Once the basins have been delineated, shared boundary bins are grouped by edge or corner contact, and each connected group is examined for one crossing. Two populated boundary bins form a candidate crossing pair if they belong to different basins and share an edge or a corner. The crossing height of a pair is the higher of its two free energies. The pair with the lowest such height defines the crossing, and its higher-energy bin gives the position marked by a cross in Figure 1c. If a boundary region contains no eligible pair, its lowest-energy populated bin provides a fallback estimate. The output identifies the basins associated with the selected crossing.

For a crossing of height *G*_×_, the apparent barrier measured from basin A is *ΔG*_*A*_ = *G*_×_ − *G*_*A*,min_, where *G*_*A*,min_ is the lowest free energy among all populated bins in A, including its boundary bins. The height of the same crossing above the neighboring basin is measured from that basin’s own minimum. Both values refer to the sampled two-dimensional surface.

#### Step 6. Selection of representative conformations

A representative conformation is selected from the observed frames of each basin core. The first core frame serves as the initial reference for least-squares superposition. The fitted coordinates are averaged, and the new mean is fitted to the preceding mean before the next round of superposition. Fitting ends when successive means differ by less than 10^−6^ Å in root-mean-square deviation (RMSD) or after 50 rounds. After the final round, the core frame with the smallest fitted RMSD from the final mean is selected. If a basin contains no core frames, its full ensemble is used. The same procedure applied to frames in a selected crossing bin yields a representative conformation for that crossing.

The representative is marked in the same frame list as the other members of its ensemble. The square in Figure 1c marks the projected position of the conformation shown in Figure 1d.

## RESULTS

### Gradient flow clustering across three molecular systems

We applied gradient flow clustering to examine protein conformational changes in all-atom MD simulations of the SARS-CoV-2 spike protein and the leucine transporter LeuT. We also examined the binding conformations sampled by a small molecule interacting with the PP2A scaffold PR65.

In our previous work, we analyzed 2.25 μs of pooled protomer trajectories from of all-atom MD simulations of the spike homotrimer to examine the transition between the closed state, with the receptor-binding domain (RBD) down, and the open state, with the RBD up. By inspecting the free-energy surface constructed from the first two PCs and examining the corresponding structures, we distinguished three main states, including a semi-open intermediate.^5^ We took a similar approach in our earlier study of LeuT, constructing a projected free-energy surface from 38.5 μs of dimer simulations. Structural analysis distinguished several substates of the outward- and inward-facing transporter, including intermediates in which access to the substrate-binding site was occluded from both sides of the membrane.^6^ For PR65, we analyzed 2.112 μs of all-atom MD simulations to characterize the binding conformations of ATUX-8385. We aligned the trajectory frames using the PR65 helices that contact the ligand and tracked the motion of ATUX-8385 relative to the protein. By inspecting the conformational map projected onto the first two PCs and examining the corresponding structures, we identified three binding poses at site S2. In separate simulations, ATUX-8385 sampled two additional binding sites.^8^ In all these studies, assigning states and substates required extensive expert inspection of projected landscapes and structural analysis of the trajectories.

Here we automate the identification of conformational states and substates directly from MD trajectory data, with no manual intervention required during state assignment. The method generates a projected free-energy surface and identifies its basins without a prescribed count. Furthermore, the automatic grid-selection rule uses the number of sampled frames and the normalized spread of conformations along each PC to estimate the grid dimensions, negating a manual input there. For each state or substate, it returns the full conformational ensemble and a representative structure from the original trajectories. It also reports the locations of candidate transition states on the surface and their representative conformations. The analyses below used 30 × 30 bins with two padding bins on each side for LeuT and ATUX-8385, and a fixed 40 × 27 grid without padding for spike.

### LeuT illustrates the stages from a grid to conformational ensembles

The first two PCs accounted for 69.2% of the variance in the coordinates of 497 LeuT C_α_ atoms. The 15,422 frames from 24 protomer trajectories formed several populated patches on the PC1– PC2 plane (Figure 2a,b). Interpolation smoothed the sampled landscape, and the intensity threshold restricted the starting pixels to the lower-energy regions (Figure 2c,d). Gradient paths from the same patch could converge near different minima. Grouping their endpoints and transferring the labels back to the original grid yielded 15 local regions (Figure 2e,f). Applying the boundary-energy and core separation criteria yielded seven global basins (Figure 2g). Excluding three shared boundary regions defined the basin cores. We then located a crossing in each boundary region and selected representative conformations (Figure 2h,i).

Basin 4 contained 7,203 frames from 17 trajectories, including frames from all ten trajectories initiated from the outward-open structure 3TT1. Trajectories initiated from the inward-open structure 3TT3 primarily populated basins 3 and 5. Basins 6 and 7 contained 23 and 79 frames, respectively, each from a single trajectory initiated from 3F3E. Figure 1d superimposes six conformations from basin 4 on the representative selected from its 7,105 core frames. The selected bin at the crossing with basin 2 contained 39 frames. The apparent heights from the minima of basins 4 and 2 were 2.21 and 1.20 k_B_T, respectively (Table 1).

**Table 1.** Candidate interbasin crossings.

| **System** | **Basin pair at crossing** | **Boundary frames** | **Frames in crossing bin** | **Barrier from first / second basin (*k*<sub>B</sub>*T*)** |
|---|---|---|---|---|
| LeuT | 1–2 | 665 | 79 | 0.00 / 0.49 |
| LeuT | 2–4 | 201 | 39 | 1.20 / 2.21 |
| LeuT | 3–5 | 278 | 37 | 1.23 / 0.85 |
| Spike | 3–4 | 1,996 | 410 | 0.75 / 1.97 |
| Spike | 1–2 | 506 | 85 | 1.34 / 2.63 |
| Spike | 2–5 | 5,806 | 620 | 0.65 / 0.64 |
| ATUX-8385 | 1–2 | 190 | 31 | 1.80 / 2.61 |

*Boundary counts include all frames in the connected region of shared bins. The spike boundary region containing the 2–5 crossing also includes bins from basin 3. Each height is measured from the minimum of the corresponding basin.*

### The spike landscape resolves five conformational ensembles

For spike, we calculated the principal components directly from the MD trajectories using the coordinates of 466 selected C_α_ atoms per protomer. PC1 and PC2 accounted for 82.8% and 4.7% of the variance, respectively. The analysis used 101,941 simulation records, including 2,000 repeated records retained in the supplied input. We projected the closed and open experimental reference structures onto these axes after fitting and included them in the map. They fell in basins 4 and 2, respectively (Figure 3a). The fixed grid used 10 Å bins with edges spanning −180 to 220 Å on PC1 and −105 to 165 Å on PC2.

**Figure 3.**
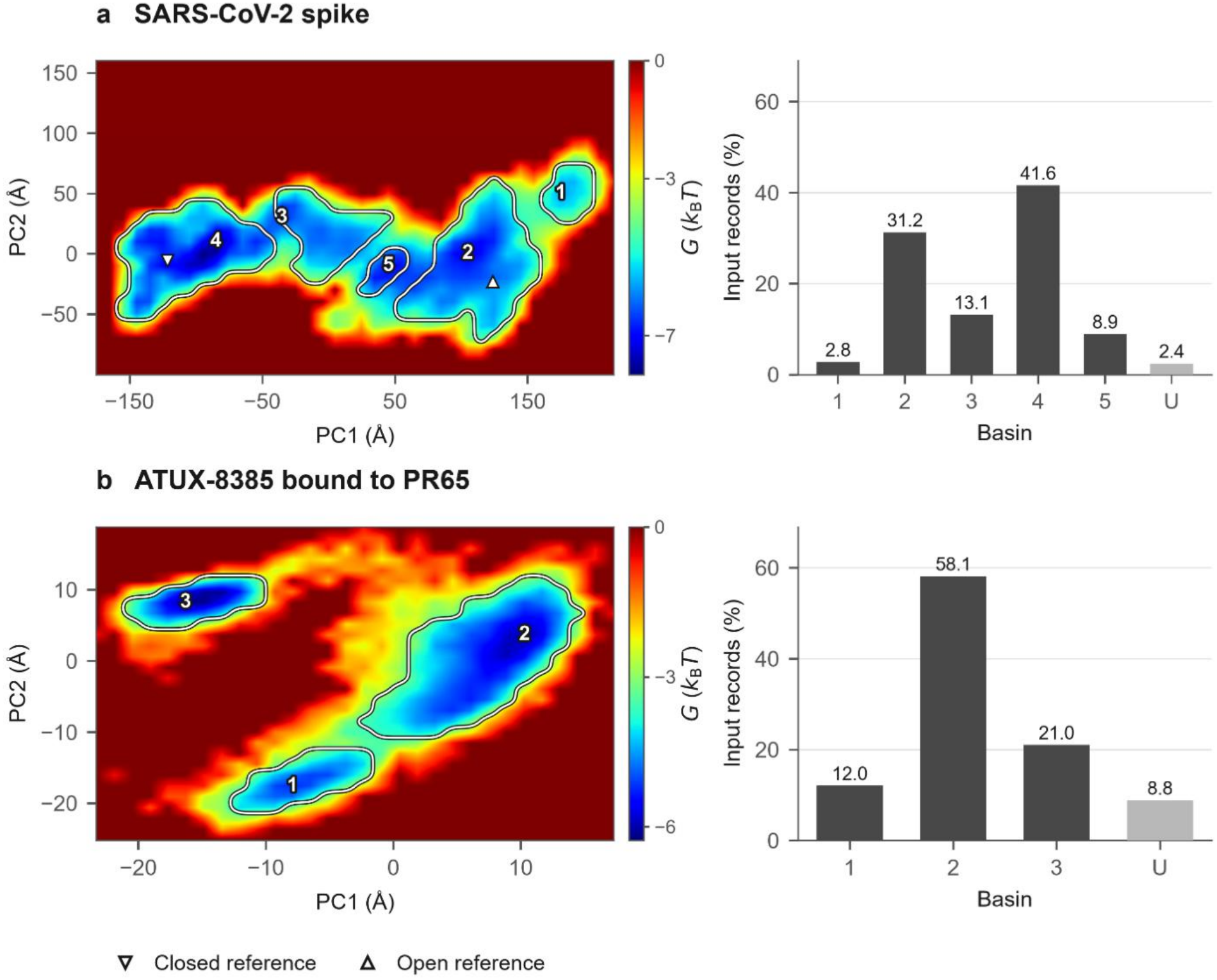
Conformational ensembles for spike and ATUX-8385. (a) Spike and (b) ATUX-8385 bound to PR65. Numbered markers identify the minima within the basin cores, and outlines delineate the cores. Triangles mark the closed and open spike reference structures. The spike PC2 sign is reversed for display. Bars give the percentage of input records assigned to each basin, including its boundary frames. U denotes unassigned records. Empty bins share the zero-energy display color.

Two basins lay between the closed and open references along PC1. The core separation criterion retained basin 3 as a separate ensemble of 13,385 frames. At the proposed merge with the closed-side region, both cores contained resolved minima and their Mahalanobis distance was 4.30. Basin 5 contained 9,039 frames, including 7,435 from one open-start conventional trajectory. Basin 1 lay beyond the open reference along PC1 and contained 2,808 frames, 2,746 of them from a single open-start steered trajectory. The crossing between basins 3 and 4 had apparent heights of 0.75 and 1.97 kBT from their respective minima (Table 1).

### ATUX-8385 occupies three ensembles at the PR65 binding site

The projected coordinates describe motion of ATUX-8385 relative to the binding site. Across three trajectories of 7,041 frames each, the first two PCs accounted for 70.9% of the variance in the coordinates of the 37 ligand atoms. Basin 2 contained 12,282 frames and was visited in all three trajectories (Figure 3b). Basin 1 contained 2,535 frames, and all 4,444 frames in basin 3 came from the third trajectory. The map placed a crossing between basins 1 and 2, with apparent heights of 1.80 and 2.61 kBT, respectively (Table 1).

The three ensembles describe alternative conformations of ATUX-8385 at site S2. The earlier structural analysis also found multiple poses at this site.^8^ The frame assignments identify the sets of structures for comparing individual basins with those poses in the receptor frame.

## DISCUSSION

A minimum on a projected free-energy surface marks a location. Structural interpretation requires knowing which conformations surround it, how much they vary, and which changes distinguish them from neighboring ensembles. Gradient flow clustering connects these levels of description. A basin can be examined as a region on the surface, as a list of frames drawn from one or more trajectories, or through a representative conformation selected from its core. The LeuT example illustrates this point. Its largest basin contains thousands of conformations sampled from several starting conditions, yet each member can be recovered as readily as the single structure shown in Figure 1d.

The choice of coordinates determines which conformational changes enter the analysis. For spike and LeuT, the selected C_α_ coordinates describe protein rearrangements. For ATUX-8385, projecting ligand coordinates after receptor alignment retains changes in both ligand pose and shape. In each case, the frames assigned to a basin allow its structural variability to be examined directly.

The spike example shows how the two merging criteria work together. The boundary-energy criterion made the two regions eligible for merging because the mean free energy along their shared boundary was slightly lower than the mean within the shallower region. Because both cores contained resolved minima and their projected ensembles were well separated relative to their spread, the core separation criterion retained the two ensembles.

Finally, occupancy has a thermodynamic interpretation when the trajectories sample the relevant equilibrium distribution. The present spike data combine conventional and steered simulations,and the LeuT data pool several substrate and ion conditions. Their crossing heights describe the sampled projections. The frame selected at a crossing provides a candidate structure for further investigation. Establishing a kinetic transition state would require dynamical evidence, such as trajectories initiated from the candidate region and followed into the adjoining states.

## CONCLUSIONS

Gradient flow clustering turns a projected free-energy landscape into explicit conformational ensembles whose members can be retrieved directly from MD trajectories. Applied to protein rearrangements and ligand binding, it connected each basin to the structures that define it and identified representative conformations at shared boundaries.

## AUTHOR CONTRIBUTIONS

Mert Gur conceived the study, designed the clustering method, led its development and evaluation, and wrote the original manuscript. Bogdan Kochetov and Shikhar Uttam jointly developed the image-segmentation component in consultation with Mert Gur, and Kochetov implemented it with guidance from Uttam. Mert Golcuk helped construct the projected free-energy surfaces and adapt the segmentation approach to them. Ethan M. Bintrim integrated the analysis scripts into a runnable pipeline. Fareeda E. Abu-Juam contributed to the projected free-energy analysis.

## CODE AND DATA AVAILABILITY

The clustering method is implemented in Python. The Gur Laboratory maintains the Python implementation. Scripts for the analyses reported here can be requested from the corresponding author. The laboratory’s GitHub profile is https://github.com/Gur-Laboratory/FELT.

The source simulations are described in references [5–8]. Requests for the trajectory data and the frame assignments used in this study may be directed to the corresponding author.

## References

1 Dror, R. O., Dirks, R. M., Grossman, J. P., Xu, H. & Shaw, D. E. Biomolecular simulation: a computational microscope for molecular biology. Annu Rev Biophys 41, 429–452 (2012). [DOI]: 10.1146/annurev-biophys-042910-155245

2 Amadei, A., Linssen, A. B. & Berendsen, H. J. Essential dynamics of proteins. Proteins 17, 412–425 (1993). [DOI]: 10.1002/prot.340170408

3 Gur, M., Zomot, E. & Bahar, I. Global motions exhibited by proteins in micro-to milliseconds simulations concur with anisotropic network model predictions. J Chem Phys 139, 121912 (2013). [DOI]: 10.1063/1.4816375

4 Li, G. et al. 3D cell nuclei segmentation based on gradient flow tracking. BMC Cell Biol 8, 40 (2007). [DOI]: 10.1186/1471-2121-8-40

5 Gur, M. et al. Conformational transition of SARS-CoV-2 spike glycoprotein between its closed and open states. J Chem Phys 153, 075101 (2020). [DOI]: 10.1063/5.0011141

6 Gur, M., Zomot, E., Cheng, M. H. & Bahar, I. Energy landscape of LeuT from molecular simulations. J Chem Phys 143, 243134 (2015). [DOI]: 10.1063/1.4936133

7 Yang-Schulz, A. et al. Direct observation of small molecule activator binding to single PR65 protein. NPJ Biosens 2, 2 (2025). [DOI]: 10.1038/s44328-024-00018-7

8 Yilmaz, S. Z. et al. Small Molecule Activator of Phosphatase PP2A Remodels Scaffold PR65 Structural Dynamics To Promote Holoenzyme Assembly. JACS Au 6, 1102–1117 (2026). [DOI]: 10.1021/jacsau.5c01514

